# Reduced heparan sulfate levels in cerebrospinal fluid reflect brain neuron correction in Sanfilippo B mice

**DOI:** 10.1101/2025.04.30.651515

**Authors:** Steven Q. Le, Alexander Sorensen, Soila Sukupolvi, Gianna Jewhurst, Grant Austin, Balraj Doray, Jonathan D. Cooper, Patricia I. Dickson

**Author notes:** Correspondence to: Patricia Dickson, MD, 4444 Forest Park Ave, Ste 5400, St. Louis, MO 63108.

## Abstract

Sanfilippo B syndrome is a rare, inherited disorder of heparan sulfate catabolism. Cerebrospinal fluid heparan sulfate (HS) has been pursued as a biomarker for CNS disease. To determine whether CSF HS reflects disease in the brain, we generated a fusion protein of the NAGLU gene encoding alpha-N-acetylglucosaminidase (the enzyme deficient in Sanfilippo B) and the transmembrane region and cytosolic tail of LAMP1, which encodes the lysosomal-associated membrane protein-1, to tether the NAGLU enzyme to the membrane and prevent cross-correction. We used AAV7 to deliver the fusion construct intravenously to Sanfilippo B mice and showed that treatment of systemic bodily organs without treatment of brain did not result in reduction of CSF HS compared with untreated Sanfilippo B mice. Next, we used intracerebroventricular delivery of the construct in AAV9 under a synapsin-1 promotor to largely confine expression to brain neurons. We showed that treatment of the brain without treatment of systemic bodily organs resulted in reduction of HS in brain and CSF. These results support the utility of CSF HS as a therapeutic biomarker for Sanfilippo B and other disorders of heparan sulfate catabolism.

---

Mucopolysaccharidoses (MPS) are a group of inherited, lysosomal diseases caused by deficiency of enzymes critical for the catabolism of glycosaminoglycans. Heparan sulfate (HS) glycosaminoglycans accumulate in several types of MPS, including Sanfilippo syndrome (MPS III), Hunter syndrome (MPS II), Hurler syndrome (MPS I), and Sly syndrome (MPS VII). Cerebrospinal fluid HS (CSF HS) has been considered as a feasible biomarker reflecting brain HS levels in MPS disorders (1). Ideally, its measurement could be used to determine the response to brain-directed therapy in clinical trials for MPS patients. Preclinical and clinical studies demonstrate a correlation of brain HS and CSF HS, suggesting a relationship (2). However, because HS is ubiquitous, it has been difficult to determine whether CSF HS truly reflects brain levels. The observation that CSF HS is lowered following intravenous enzyme replacement therapy (which largely does not cross the blood-brain barrier, BBB) suggests the possibility that HS may cross from blood into CSF (3).

Experiments designed to test this hypothesis must prevent the cross-correction that occurs with secreted, soluble lysosomal enzymes. To this end, we generated an expression construct with the *NAGLU* gene encoding alpha-N-acetylglucosaminidase (the enzyme deficient in MPS IIIB), a 6-glycine linker, and a 114-bp nucleotide segment corresponding to the transmembrane region and cytosolic tail of *LAMP1*, which encodes the lysosomal-associated membrane protein-1, followed by a c-Myc epitope tag after the cytosolic tail, following published methods (4). To test expression *in vitro*, human MPS IIIB fibroblasts were transduced with lentiviral NAGLU-LAMP1 (**Supplemental Methods**). Cells and media were harvested and assayed for NAGLU activity as described (5). We confirmed intracellular NAGLU activity but found no NAGLU activity in the secreted media, consistent with successful membrane tethering (**Supplemental Figure 1**). We then performed *in vivo* testing in *Naglu*^*-/-*^ mice. To deliver NAGLU-LAMP1 systemically but avoid treating brain disease, we used adeno-associated viral vector-7 (AAV7), which does not cross the BBB when administered intravenously. We delivered 1.5×10^11^ vector genomes per mouse of AAV7-NAGLU-LAMP1 by tail vein at four weeks of age, avoiding the neonatal period during which the BBB is not completely intact. Four weeks post-dose, NAGLU activity was detected in systemic bodily organs but not in brain (**Figure 1A**), with concomitant decrease in β-hexosaminidase (β-Hex) activity in the organs showing good NAGLU activity (**Supplemental Figure 2A**). Consistent with lack of secretion of the membrane-tethered enzyme, NAGLU activity was not detected in serum (**Figure 1A**). Brain homogenates, serum, and CSF were assayed for HS by glycan reductive isotope labeling liquid chromatography/mass spectrometry (GRIL-LC/MS) (6). Serum HS, but not brain or CSF HS, was reduced in treated *Naglu*^*-/-*^ mice compared to untreated controls (**Figure 1B-D**).

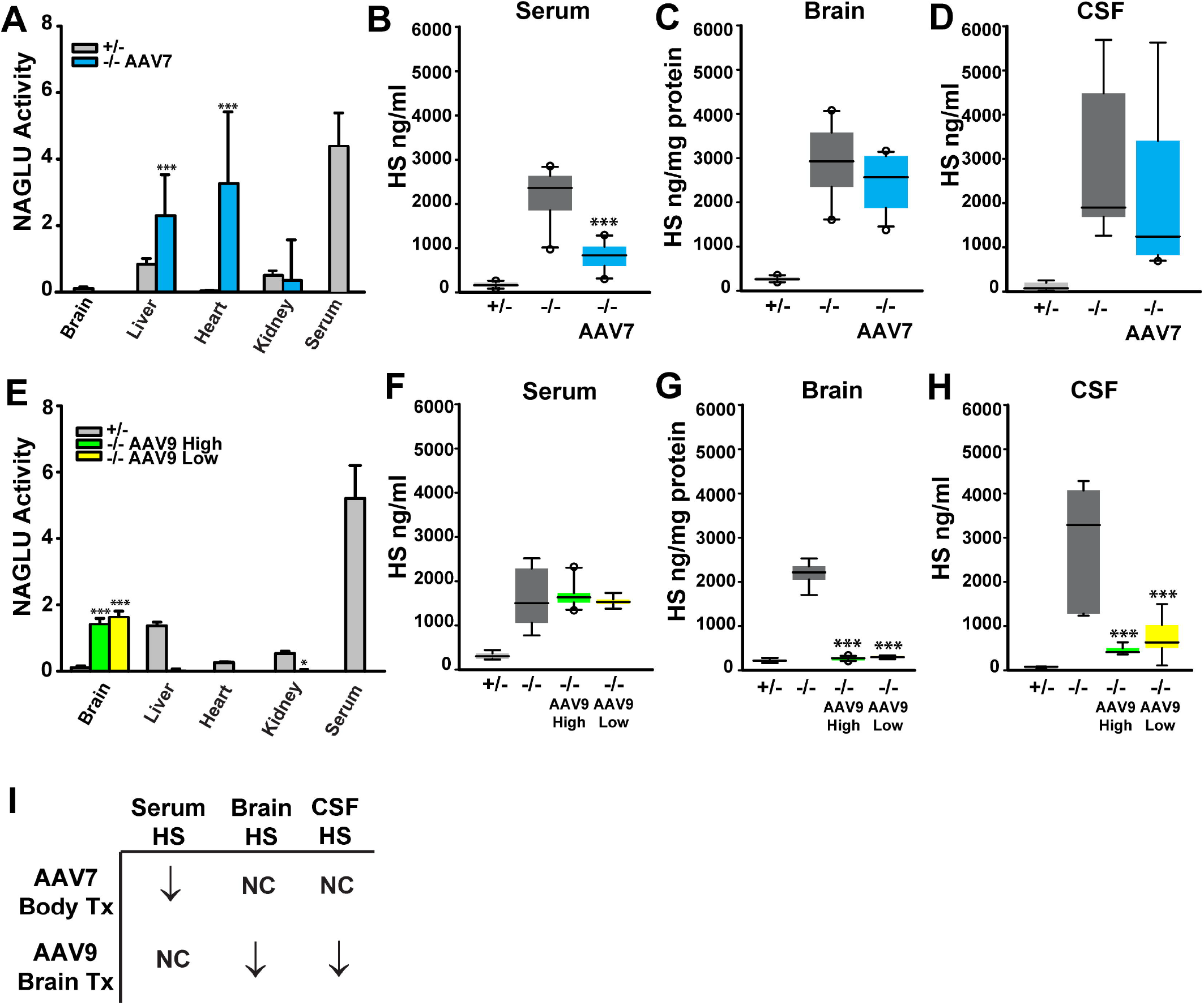
(**A**) NAGLU enzymatic activity in 8 wk old *Naglu*^-/-^ mice (n=12: 8 females, 4 males) treated at 4 wk with AAV7-NAGLU-LAMP1 (“AAV7”) intravenously, compared to untreated -/- mice (n=10: 5 females, 5 males) and +/- controls (n=10: 5 females, 5 males). Mean and S.D. shown. *Naglu*^-/-^ mice have undetectable NAGLU activity (not shown). (**B-D**) Total HS in serum, CSF, and brain of mice treated with AAV7-NAGLU-LAMP1 and controls. Boxplots depict median, quartile; whiskers show 10^th^ and 90^th^ centiles, circles are 5^th^ and 95^th^ centiles. ***p<0.001 vs -/-. (**E**) NAGLU enzymatic activity in 4 wk old *Naglu*^-/-^ mice treated intracerebroventricularly at P1 or P2 with 6.5×10^10^ vector genomes (“AAV9-High;” n= 13: 3 females, 10 males) or 6.5×10^9^ vector genomes (“AAV9-Low;” n= 8: 4 females, 4 males) AAV9-Syn1-NAGLU-LAMP1, compared to untreated -/- mice (n=8: 2 females, 6 males) and +/- controls (n=7: 3 females, 4 males). Mean and S.D. shown. *Naglu*^-/-^ mice have undetectable NAGLU activity (not shown). (**F-H**) Total HS in serum, CSF, and brain of the mice treated with AAV9-Syn1-NAGLU-LAMP1 and controls. Boxplots depict median, quartile; whiskers show 10^th^ and 90^th^ centiles. *p<0.05 and ***p<0.001 vs -/-. (**I**) Summary of experimental results. Tx: treatment. Values for all data points in graphs are reported in the Supporting Data Values file.

Next, we designed a separate experiment to study the effect on CSF HS when NAGLU restoration is largely confined to brain neurons. For this, we used an AAV9 vector with NAGLU-LAMP1 expression under the control of a synapsin-1 promoter. We administered 6.5×10^10^ vector genomes (“high dose”) or 6.5×10^9^ vector genomes (“low dose”) per mouse of AAV9-Syn1-NAGLU-LAMP1 into one lateral cerebral ventricle of *Naglu*^-/-^ mice at P1 or P2. Four weeks post-dose, NAGLU activity was readily detected in brains of mice treated with both vector doses, while NAGLU activity in the periphery including heart, kidney, liver, and serum was undetectable (low dose) or extremely low (high dose) (**Figure 1E**). As before, β-Hex activity was only significantly decreased in the organs expressing NAGLU (**Supplemental Figure 2B**). Confocal microscopy confirmed neuronal expression and distribution of NAGLU-LAMP1 (**Supplemental Figure 3**). We also observed a decrease in immunostaining for astrocytes and activated microglia in the brains of treated *Naglu*^-/-^ mice, supporting a therapeutic effect (**Supplemental Figure 4**). GRIL-LC/MS showed that brain HS was normalized and CSF HS was reduced in the mice that were treated intracerebroventricularly with AAV9-Syn1-NAGLU-LAMP1, whereas there was no difference in serum HS between treated and untreated *Naglu*^*-/-*^ mice (**Figure 1F-H**).

These results, summarized in **Figure 1I**, suggest that CSF HS reflects HS in the central nervous system. Moreover, as CSF HS was normalized with restoration of NAGLU activity in brain neurons, the results support the utility of CSF HS as a therapeutic biomarker for gauging improvement in brain disease due to MPS.

## Supporting information

raw data file

## Notes

Conflicts of Interest. J.D.C. receives research support from Neurogene, Regenexbio and Alnylam and is a consultant with JCR pharmaceuticals. P.I.D. receives research support from Alnylam, BioMarin Pharmaceutical Inc., and M6P Therapeutics and is an inventor on Patent #USSN 15/946,505 Enzyme replacement therapy for mucopolysaccharidosis IIID. B.D. is an inventor on Patent #US 10,907,139 B2 Compositions comprising a modified GlcNAc-1-phosphotransferase and methods of use thereof.

### Competing Interest Statement

J.D.C. receives research support from Neurogene, Regenexbio and Alnylam and is a consultant with JCR pharmaceuticals. P.I.D. receives research support from Alnylam, BioMarin Pharmaceutical Inc., and M6P Therapeutics and is an inventor on Patent #USSN 15/946,505 Enzyme replacement therapy for mucopolysaccharidosis IIID. B.D. is an inventor on Patent #US 10,907,139 B2 Compositions comprising a modified GlcNAc-1-phosphotransferase and methods of use thereof.

### Funding

- National Institute of Neurological Disorders and Stroke: RM1 NS132962

