## Supplemental materials for "Reduced heparan sulfate levels in cerebrospinal fluid reflect brain neuron correction in Sanfilippo B mice"

Reduced levels of heparan sulfate in cerebrospinal fluid reflect correction of brain neurons in mucopolysaccharidosis type III mice

Steven Q. Le, Alexander Sorensen, Soila Sukupolvi, Gianna Jewhurst, Grant Austin, Balraj Doray, Jonathan D. Cooper, Patricia I. Dickson

Department of Pediatrics, Washington University School of Medicine, St. Louis, MO

**Supplemental Material**

### Methods

**Sex as a biological variable**. We studied male and female mice as reported in the Figure Legend. We did not observe sex-specific differences in the outcomes we assessed.

**Reagents**. Human MPS type IIIB fibroblasts (GM 1426) were obtained from Coriell. 4-methylumbelliferyl-*N*-acetyl-α-glucosaminide and 4-methylumbelliferyl-*N*-acetyl-β-glucosaminide were obtained from EMD Millipore Chemicals (Darmstadt, Germany). TrueBlack Lipofuscin Autofluorescence Quencher was obtained from Biotium. Primary antibodies used in this study were rabbit anti-NAGLU (ab214671, abcam), mouse anti-NeuN (MAB377, Sigma), anti-mouse GFAP (G3893, Sigma), and anti-rat CD68 (MCA1957, Biorad). Secondary antibodies purchased from Thermofisher were goat-anti rabbit 488 (A11008), goat anti-mouse 546 (A-11003), and goat anti-rat 647(A-21247).

**Viral vectors**. The AAV vectors used in this study were constructed either by inserting the transgene NAGLU-LAMP1 into an AAV-CBA backbone containing the cytomegalovirus (CMV) enhancer/chicken beta actin (CBA) promoter, or an AAV vector driven by the neuron-specific synapsin 1 promotor (Syn1). The NAGLU-LAMP1 transgene construct consists of the human NAGLU cDNA joined via a linker nucleotide sequence (encoding six glycine residues) to a 114-bp nucleotide segment corresponding to the transmembrane region and cytosolic tail of LAMP1. Nucleotides encoding the c-Myc epitope tag were appended to the cytosolic tail of LAMP1. The NAGLU-LAMP1 transgene was initially cloned into a commercially available lentiviral vector, Lenti-III-pGK (G305, abm) to generate LV-NAGLU-LAMP1to test transduction *in vitro*. For mouse studies, the NAGLU-LAMP1 transgene in the AAV vectors were packaged into either the AAV7 serotype (for CBA NAGLU-LAMP1) or the AAV9 serotype (for Syn1 NAGLU-LAMP1) and produced by the Hope Center Viral Vectors Core at Washington University School of

Medicine. The AAV7-CBA-NAGLU-LAMP1-myc viral titer was 1.5x$10^{13}$ viral genomes per ml, while the AA9-Syn1-NAGLU-LAMP1-myc viral titer was 1.3x$10^{13}$ viral genomes per ml.

**Cell culture**. Human MPS IIIB fibroblasts (GM 1426) were transduced with LV-NAGLU-LAMP1 and selected with 10 µg/ml puromycin added to the culture media. Cells and media were harvested and tested for NAGLU and beta-hexosaminidase activity as described below.,

**Mice**. All experiments in this study were performed using the *Naglu*$^{-/-}$ mouse strain B6.129S6-Naglu(tm1Efn)/J . Mice were bred by crossing knockout males to heterozygous females, producing 50% knockout and 50% heterozygous offspring. Genotyping was performed at birth with primer set NAG5′: TGGACCTGTTTGCTGAAAGC and NAG3′: CAGGCCATCAAATCTGGTAC for *Naglu* wild-type alleles, and primer set Neo5′: TGGGATCGGCCATTGAACAA; Neo3′: CCTTGAGCCTGGCGAACAGT for *Neo*$^{r}$ mutant alleles. Intravenous dosing of AAV7-NAGLU-LAMP1 was performed by tail vein injection. Intracerebroventricular dosing of AAV9-Syn1-NAGLU-LAMP1 was performed by intracerebroventricular injection in neonatal mice as described (1). For intraventricular dosing of AAV9-Syn1-NAGLU-LAMP1, mice were cryoanesthetized, and viral vector was injected at 5 µl volume using a 32-gauge Hamilton syringe. Terminal CSF collection was performed by cisterna magna puncture. Blood was obtained by cardiac puncture, following which mice were perfused transcardially with chilled PBS. Brain left hemisphere, liver, kidneys, and heart were collected and snap frozen for biochemical assays. The brain right hemisphere was fixed in PFA for histology.

**Enzymatic activity assays**. Samples were coded so that assays could be performed in a blinded fashion. NAGLU activity and total hexosaminidase activity were quantified using 4-methylumbelliferyl assays as previously described (1). Briefly the enzymatic activity of NAGLU was determined by hydrolysis of the fluorogenic substrate, 4-methylumbelliferyl-*N*-acetyl-α-glucosaminide, obtained from EMD Millipore Chemicals (Darmstadt, Germany) with minor modifications of a published protocol, using 0.1 mM substrate in the incubation mixture (2). Enzymatic activity of beta-hexosaminidase (combined A and B isoforms) was determined by hydrolysis of 4-methylumbelliferyl-*N*-acetyl-β-glucosaminide (EMD Millipore Chemicals, Darmstadt, Germany) using 1.25 mM substrate in the incubation mixture. For both enzymes, a unit of activity is defined as release of 1 nmol of 4-methylumbelliferone (4MU) per hour. Protein concentration was estimated by the Bradford method, using bovine serum albumin as a standard.

**HS quantification**. Samples were sent to the University of California San Diego GlycoAnalytics core for evaluation of HS levels by mass spectrometry. Samples were coded so that assays were performed in a blinded fashion. For HS analysis, tissue homogenates, CSF, and serum samples were treated with protease and the negatively charged glycosaminoglycans were trapped using Anion Exchange Chromatography (AEC). Bound glycosaminoglycans were eluted with high concentration of chloride as counter ion (2M NaCl solution) followed by desalting using pre-packed desalting column (PD-10). Purified total glycosaminoglycans were digested with mixture of Heparinase (I, II & III) which depolymerize the chains, generating mostly disaccharides. The disaccharides were tagged with $^{12}C_6$-aniline by reductive coupling method (termed as Glycan Reductive Isotopic Labeling, GRIL)(3). These aniline-tagged disaccharide mixtures were spiked with known amounts of $^{13}C_6$-isotopic aniline tagged disaccharide standard mixture and analyzed using Liquid Chromatography Mass Spectrometry (LC-MS) in negative ionization mode (LTQ-Orbitrap Discovery, Thermo Scientific). The ion intensities of the standards and the samples that coelute were deconvoluted by mass spectrometry and quantified.

**Immunostaining**. Brains were post-fixed overnight in PFA and cryoprotected in 30% sucrose/50 mM tris-buffered saline (TBS) at 4 ºC. Samples were coded so that histology was performed in a blinded fashion. A 1-in-6 series of 40 µm coronal forebrain hemisections from each mouse were collected in 96-well plates containing a cryoprotectant solution (TBS/30% ethylene glycol/15% sucrose/0.05% sodium azide). Sections were subsequently mounted and stained immunofluorescently stained using 1° antibody and 2° antiserum in TBST with 10% normal serum as described previously, counterstaining with the TrueBlack Lipofuscin Autofluorescence Quencher (Biotium) (4-6). Primary antibodies included rabbit anti-NAGLU (ab 214671), mouse anti-NeuN (MAB377), mouse anti-GFAP (G3893), and rat-CD68 (MCA 1957), with corresponding secondary antibodies goat anti rabbit 488 (A11008), goat anti-mouse 546 (A11003) and goat anti-rat 647 (A21247). Stained slides were scanned using a Zeiss Axio Imager.Z1 microscope and processed with the StereoInvestigator (MBF Bioscience) software, and quantification of immunoreactivity for each antigen was performed via thresholding using ImagePro Premier as described (4-6).

**Statistical analysis**. We used STATA 17 (College Station, Texas) to perform analysis of variance for categorical variables using the “anova” command. We performed post-hoc pairwise comparisons using the “pwmean” command.

**Study approval**. All study procedures involving mice were in full compliance with the National Institutes of Health's Guide for the Care and Use of Laboratory Animals and approved by the Animal Studies Committee in the Division of Comparative Medicine at Washington University School of Medicine. Laboratory procedures including viral vectors were approved by the Biosafety Committee at Washington University School of Medicine. Human fibroblasts were obtained from Coriell as a deidentified cell line under a Materials Transfer Agreement.

**Data availability**. All data shown in the figures are provided in a Supporting Data Values file (Microsoft Excel format).

**Acknowledgements**. Support was provided by 1RM1 NS132962 to P.I.D. and J.D.C. and by the Hope Center Viral Vectors Core at Washington University School of Medicine.

**Supplemental Figure Legends**

**Supplemental Figure 1**. *In vitro* expression of NAGLU-LAMP1 in human fibroblasts. (A) Schematic of construct. Human NAGLU cDNA, 6 glycine spacer (6G), transmembrane domain of lysosomal-associated membrane protein-1 (LAMP1), and c-myc, flanked by restriction sites SalI and XhoI. (**B**) GM 1426 cells (human MPS IIIB fibroblasts) were transduced in 24-well plates with lentiviral (LV) NAGLU-LAMP1. After washing the cells, media and cell pellets were each assayed. NAGLU activity represents mean and s.d. of 2 experiments and demonstrates that the construct is enzymatically active and not secreted. Beta-hexosaminidase activity in GM 1426 cells and media suggesting the construct is able to catabolize lysosomal substrate.

**Supplemental Figure 2**. Beta hexosaminidase activity in treated *Naglu*-/- mice and controls. (**A**) Enzymatic activity of total beta-hexosaminidase ("β-Hex") in organs of 8 wk old *Naglu*-/- mice (n=12: 8 females, 4 males) treated at 4 wk of age with AAV7-NAGLU-LAMP1 ("AAV7") intravenously, compared to untreated -/- (n=10: 5 females, 5 males) and +/- (n=10: 5 females, 5 males) mice. (**B**) Total beta-hexosaminidase activity in organs of 4 wk old *Naglu*-/- mice treated intracerebroventricularly at P1 or P2 with $6.5 \times 10^9$ vector genomes AAV9-Syn1-NAGLU-LAMP1 ("AAV9-Low;" n=8: 4 females, 4 males) or $6.5 \times 10^{10}$ vector genomes AAV9-Syn1-NAGLU-LAMP1 ("AAV9-High;" n=13: 3 females, 10 males), compared to untreated -/- (n=8: 2 females, 6 males) and +/- (n=7: 3 females, 4 males) mice. Means and S.D. shown. **$p<0.01$ and ***$p<0.001$ vs -/-.

**Supplemental Figure 3**: Immunofluorescence staining demonstrating cellular and tissue localization of NAGLU following intracerebral injection of $6.5 \times 10^{10}$ vector genomes (high dose) AAV9-Syn1-NAGLU-LAMP1. (**A**) Anti-NAGLU staining (green) of a coronal hemispheric section of brain neocortex. Scalebar: 1000 µm. (**B**) Absence of colocalization of NAGLU staining (green) with staining for CD68 (cyan, arrows) or GFAP (red, arrowheads). Scalebar: 50 μm. (**C**) Cellular colocalization of NAGLU staining (green) with neuronal nuclei (red). Scalebar: 100 μm.

**Supplemental Figure 4**. Quantitative immunofluorescence staining for activated microglia (CD68) and astrocytosis (GFAP) in 4 wk old *Naglu*-/- mice treated at P1 or P2 with $6.5x10^{10}$ vector genomes AAV9-Syn1-NAGLU-LAMP1 (“AAV9-High,” n=5: 2 females, 3 males) intracerebroventricularly, compared to untreated -/- (n=4: 1 female, 3 males) and +/- (n=4: 2 females, 2 males) mice. (A) Immunofluorescence staining for CD68 or GFAP at 10x magnification. Scalebar: 50 μm. (B) Quantification of immunofluorescence staining. Bars represent means and S.D. of immunofluorescence intensity above threshold as a percentage of the area evaluated.

**A**

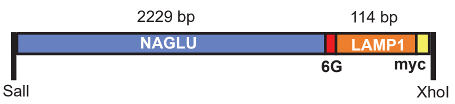


**B**

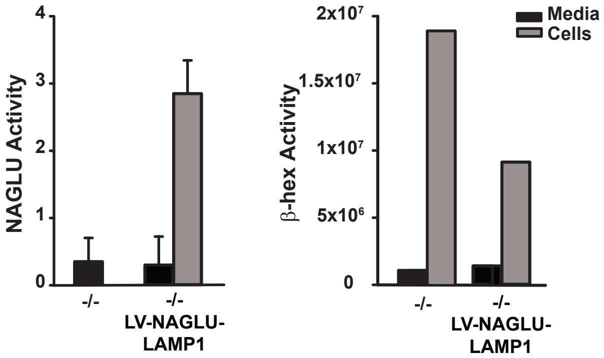


Supplemental Figure 1

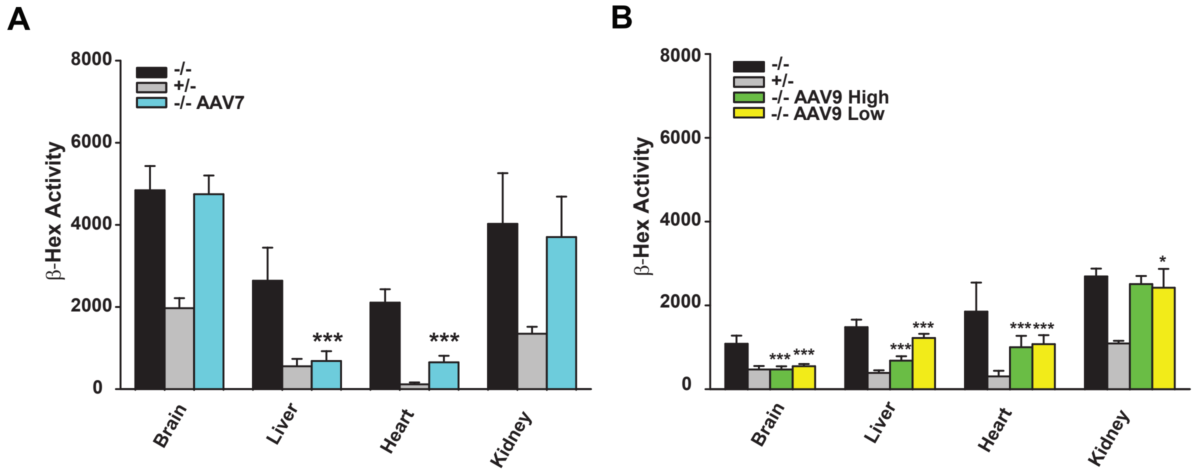


Supplemental Figure 2

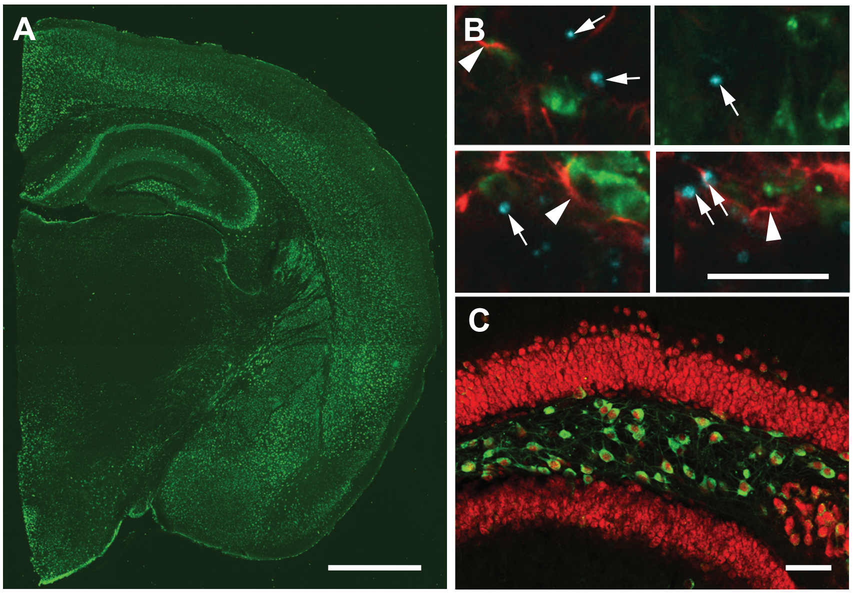


Supplemental Figure 3

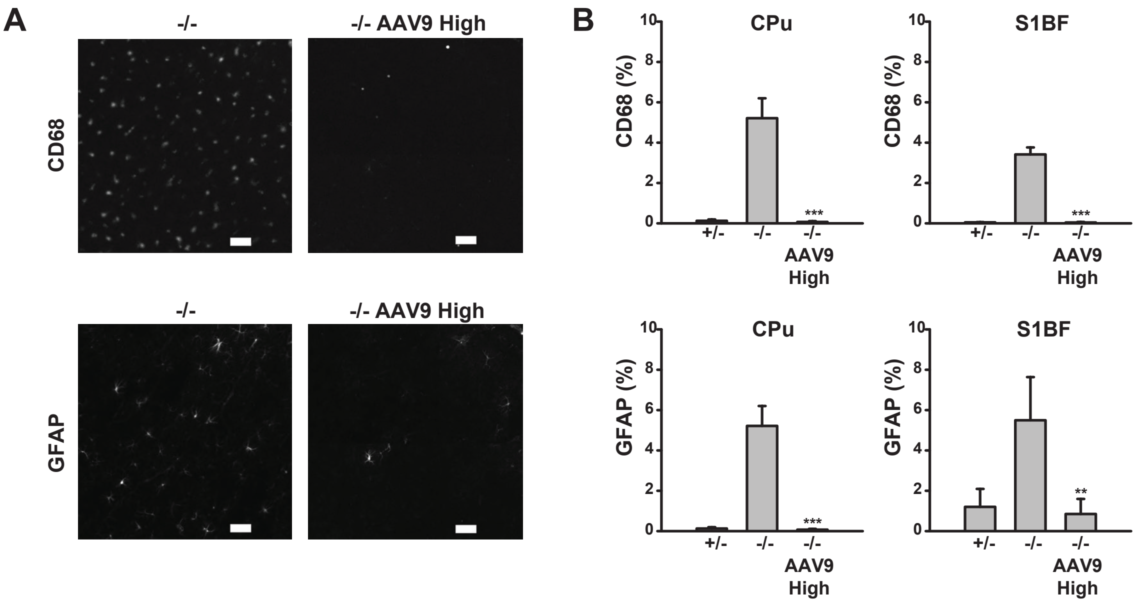


Supplemental Figure 4
